# Hippocampal CA1 neurons are crucial for sleep-associated memory formation in humans: The role of theta power during NREM sleep

**DOI:** 10.64898/2026.09.02.748785

**Authors:** A. Hanert, E.M. Kurz, F.D. Weber, J. Döhring, A. Pedersen, A. Burgalossi, J. Born, T. Bartsch

**Author notes:** Corresponding author: Thorsten Bartsch, MD, Memory Disorders and Plasticity Group Dept. of Neurology, University Hospital Schleswig-Holstein, Kiel Arnold Heller Str. 3, 24105 Kiel, Germany. equally contributing (A. Hanert, E.M. Kurz).

## Abstract

The formation of long-term memory during sleep depends on the reactivation and redistribution of recently acquired mnemonic information during non-rapid eye movement (NREM) sleep. Animal studies suggest that hippocampal memory replay during slow-wave sleep is coordinated through the interaction of sharp-wave ripples, thalamocortical sleep spindles, and neocortical slow oscillations (SOs). However, direct evidence for the contribution of hippocampal network dynamics to sleep-dependent memory consolidation in humans remains limited.

Here, we investigated sleep-dependent memory consolidation in patients (n=13) with transient global amnesia (TGA), a clinical syndrome associated with focal and transient lesions of the hippocampal CA1 region. Patients completed a verbal paired-associative learning task followed by nocturnal polysomnography and subsequent memory retrieval during the acute phase of TGA (acute condition) and again after clinical recovery (follow-up condition).

Overnight memory consolidation was significantly impaired during the acute phase compared with the follow-up session. NREM EEG theta power (4-8 Hz) was reduced during the acute phase of TGA. Importantly, increases in theta power from the acute to the follow-up session predicted corresponding improvements in memory consolidation within individuals. In contrast, established NREM markers of sleep-dependent memory consolidation, including sleep spindle density, SO density, and SO-spindle coupling, did not differ between the acute and follow-up conditions.

These findings suggest that transient hippocampal CA1 dysfunction disrupts sleep-related hippocampal network dynamics reflected in reduced NREM theta activity, which in turn is associated with impaired memory consolidation. Sleep-related theta oscillations may therefore represent a functional marker of hippocampal network integrity during sleep-dependent memory consolidation in humans.

## Introduction

Consider the experience of struggling to remember newly learned information, such as unfamiliar name-face associations or foreign vocabulary, only to find it more readily retrievable after a night of sleep. Such improvements are thought to reflect more than the mere passage of time. During sleep, the brain reactivates recently encoded experiences, allowing fragile memory traces to stabilize, reorganize, and gradually integrate into long-term memory networks [1]. A central question is how this neural reactivation is coordinated in the human brain. This process is particularly relevant for episodic memories, which initially depend on the hippocampus and are thought to become increasingly represented by distributed neocortical networks over time [1,2]. During slow-wave sleep, newly encoded memory representations are gradually redistributed from hippocampal short-term to long-term storage sites in neocortical networks, according to the Standard Model of Systems Consolidation [3–6]. This process is thought to rely on the repeated reactivation of hippocampal firing patterns that were encoded during prior waking states [7,8].

During NREM sleep, memory reactivation and hippocampal-neocortical communication are thought to be supported by the precise temporal coordination of neural oscillations across hippocampal, thalamocortical, and neocortical networks. High-frequency sharp-wave ripples (∼80-150 Hz in humans), generated in hippocampal CA1 and CA3 subfields and considered critical for memory replay [9,3], are nested within thalamocortical spindle oscillations (∼15 Hz), which in turn are coordinated by neocortical slow oscillation (SO; < 1 Hz) [10–13]. This hierarchical coupling of sleep oscillations is thought to facilitate the transfer of memory representations from hippocampal to neocortical networks [1,14].

More recently, accumulating evidence suggests that theta oscillations may represent an additional mechanism supporting memory reactivation during sleep. Studies employing targeted memory reactivation in humans have demonstrated that memory reactivation during sleep is accompanied by increases in theta activity [15–18]. At the same time, theta oscillations are also a well-established feature of active wakefulness and facilitate the encoding of novel information as well as the induction of synaptic plasticity within hippocampal networks [19–21]. Notably, memory reactivation during wakefulness has been hypothesized to effectively induce synaptic plasticity when hippocampal CA1 inputs receive burst stimulation during ongoing theta oscillations [22]. Thus, theta activity may provide a unique window into hippocampal contributions to memory reactivation and synaptic plasticity.

During sleep, theta activity is not restricted to wakefulness and REM sleep but is also present during NREM sleep [23,24]. While some studies suggest that neocortical theta activity can be temporally coordinated with SOs, reflecting large-scale network interactions [25,24], theta activity more generally has been implicated in memory reactivation processes during sleep [23]. Although scalp-recorded theta activity cannot be attributed exclusively to the hippocampus, converging evidence suggests that sleep-related increases in theta activity may partly reflect hippocampal or hippocampal-neocortical network dynamics involved in memory reactivation [26,16,23]. Because CA1 plays a critical role in hippocampal output, memory replay, and synaptic plasticity [27,28], acute CA1 dysfunction would be expected to disrupt theta-related sleep activity. Although changes in other consolidation-related oscillations, such as SOs and spindles, cannot be excluded, theta activity may be particularly sensitive to transient hippocampal disruption.

To address this issue, we investigated sleep-dependent memory consolidation in patients with transient dysfunction of the CA1 region. Hippocampal CA1 lesions constitute a characteristic neural correlate of the acute stage of transient global amnesia (TGA), a rare syndrome of selective anterograde and retrograde memory loss that resolves within 24 hours [29–34]. This condition therefore provides a unique human model for studying the contribution of the hippocampal CA1 region to memory consolidation.

We hypothesized that hippocampus-dependent memory consolidation during sleep would be impaired during the acute stage of TGA and would recover following resolution of hippocampal dysfunction. We further predicted that acute CA1 dysfunction would be associated with altered theta activity during NREM sleep, consistent with a disruption of hippocampal or hippocampal-neocortical network dynamics involved in memory reactivation. By combining behavioral memory testing with polysomnographic recordings during the acute and recovery phases of TGA, we aimed to clarify how hippocampal CA1 integrity contributes to sleep-dependent memory consolidation and sleep-related oscillatory dynamics in humans.

To test these hypotheses, we assessed patients with TGA during the acute phase and again after clinical recovery. We first examined whether overnight memory consolidation in a paired-associate learning task was impaired during acute CA1 dysfunction. We then compared sleep architecture and NREM oscillatory activity across phases, focusing on theta power, SOs, spindles, and SO-spindle coupling. Finally, we tested whether recovery-related changes in sleep oscillations were associated with improvements in memory consolidation.

## Results

### Effect of sleep on word pair association

To confirm successful encoding of word pairs in both conditions, we analyzed polynomial contrasts from one-way repeated-measures ANOVAs with learning trial (1 to 5) as the within-subject factor. These analyses revealed significant linear learning trends in both conditions (acute: *p* = .003, follow-up: *p* = .002). Consistent with our previous work [35,33,36], a repeated-measures ANOVA including condition (acute vs. follow-up) and learning trial as within-subject factors revealed a significant main effect of condition (*F*(1, 13) = 30.38, *p* < .001), indicating significantly better memory recall for word pairs during follow-up session across all learning trials. Post-hoc comparisons confirmed significantly higher recall in the follow-up condition at each trial (all *p*’s < .05, Holm-Bonferroni corrected for 5 comparisons).

To assess overnight changes in memory performance, we conducted a 2 x 2 repeated measures ANOVA with condition (acute vs. follow-up) and recall (immediate vs. delayed recall) as within-subject factors. A significant condition x recall interaction was observed (*F*(1, 13) = 6.62, *p* = .023), indicating differential overnight retention between conditions. During the follow-up session, memory performance showed only a modest decline from immediate to delayed recall (89.82 ± 2.91% vs. 81.61 ± 2.38%), whereas patients in the acute phase exhibited markedly greater overnight forgetting (68.16 ± 4.90% vs. 43.27 ± 4.80%; Figure 1).

**Figure 1.**
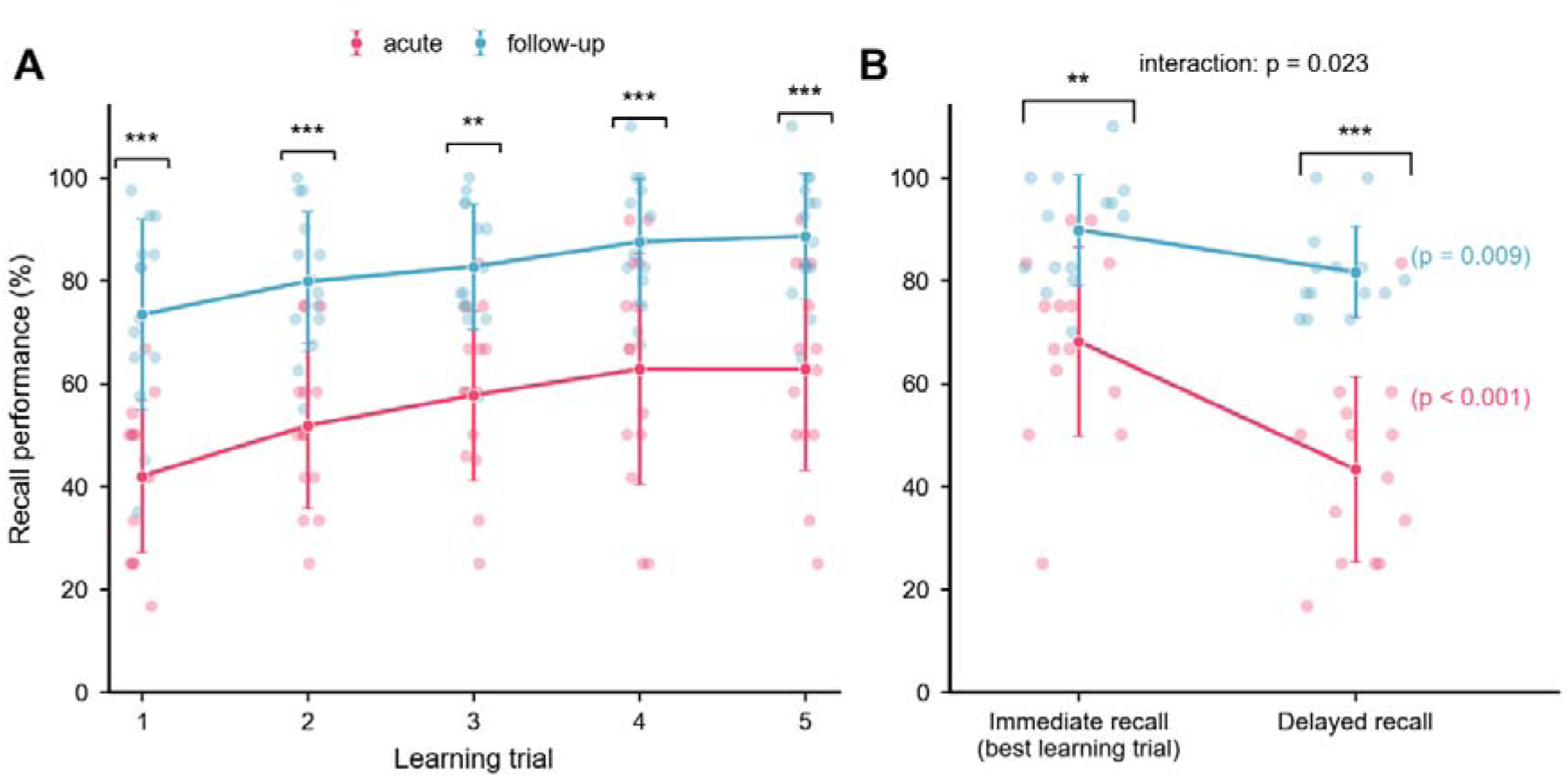
(A) Learning curves across five trials in the acute and follow-up conditions. Performance is expressed as the percentage of correctly recalled word pairs and shown as mean ± SD, with individual data points overlaid. Memory performance increased across learning trials in both conditions. Asterisks indicate significant differences between conditions at the respective trials. (B) Memory performance during immediate recall (best learning trial) and delayed recall. Data are shown as mean ± SD with individual data points. Asterisks denote significant differences between conditions, and p-values in parentheses indicate within-condition changes from immediate to delayed recall. The significant condition × recall interaction is indicated above the panel. ** *p* < .01, *** *p* < .001

### Hippocampal CA1 lesions impair memory consolidation

In 12 patients, we detected a total of 14 hippocampal lesions on axial MRI scans obtained within 24-72 hours of TGA symptom onset. Five patients showed lesions in the left hippocampus, four in the right hippocampus, and three showed bilateral lesions. Along the anterior-posterior axis of the hippocampus, lesion distribution was heterogeneous: 64.29% of all lesions were located in the middle portion of the hippocampus, whereas 21.43% and 14.29% were located in the anterior and posterior portions, respectively (Figure 2C).

**Figure 2.**
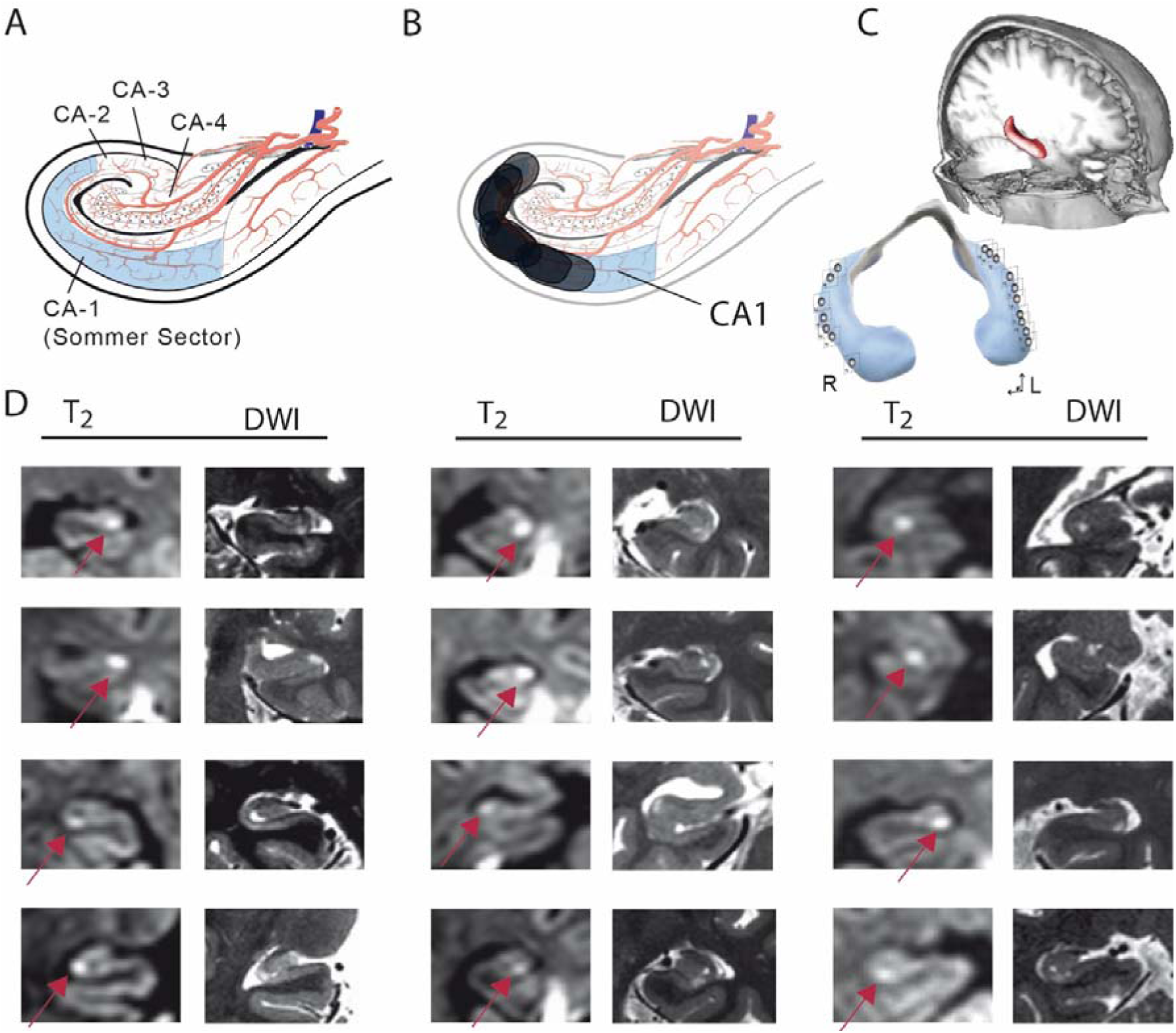
A) Schematic representation of a coronary slice of the hippocampal cornu ammonis indicating sectors after Lorente de Nó. B) Synopsis of all T2/DWI lesions mapped onto the anatomical template of the cornu ammonis. C) Three-dimensional model of the hippocampus and location within medial temporal lobe. Lesions are mapped along the anterior-posterior axis on both sides. D) MRI of representative patients shows restriction of lesions to the area CA1.

### Sleep stages and micro-architecture during CA1 lesions and after recovery

Patients with TGA spent significantly less time in bed and obtained significantly less total sleep time during the acute phase than during the follow-up night (Table 1). Specifically, time spent in N1, N2, and REM sleep was significantly reduced during the acute phase, whereas time spent in N3 did not differ significantly between conditions. Relative to time in bed, the proportion of N2 and REM sleep were significantly lower during the acute phase, while the proportion of wake after sleep onset (WASO) was significantly higher. No significant differences were observed for the relative amounts of N1 or N3 sleep (Table 1).

**Table 1.**
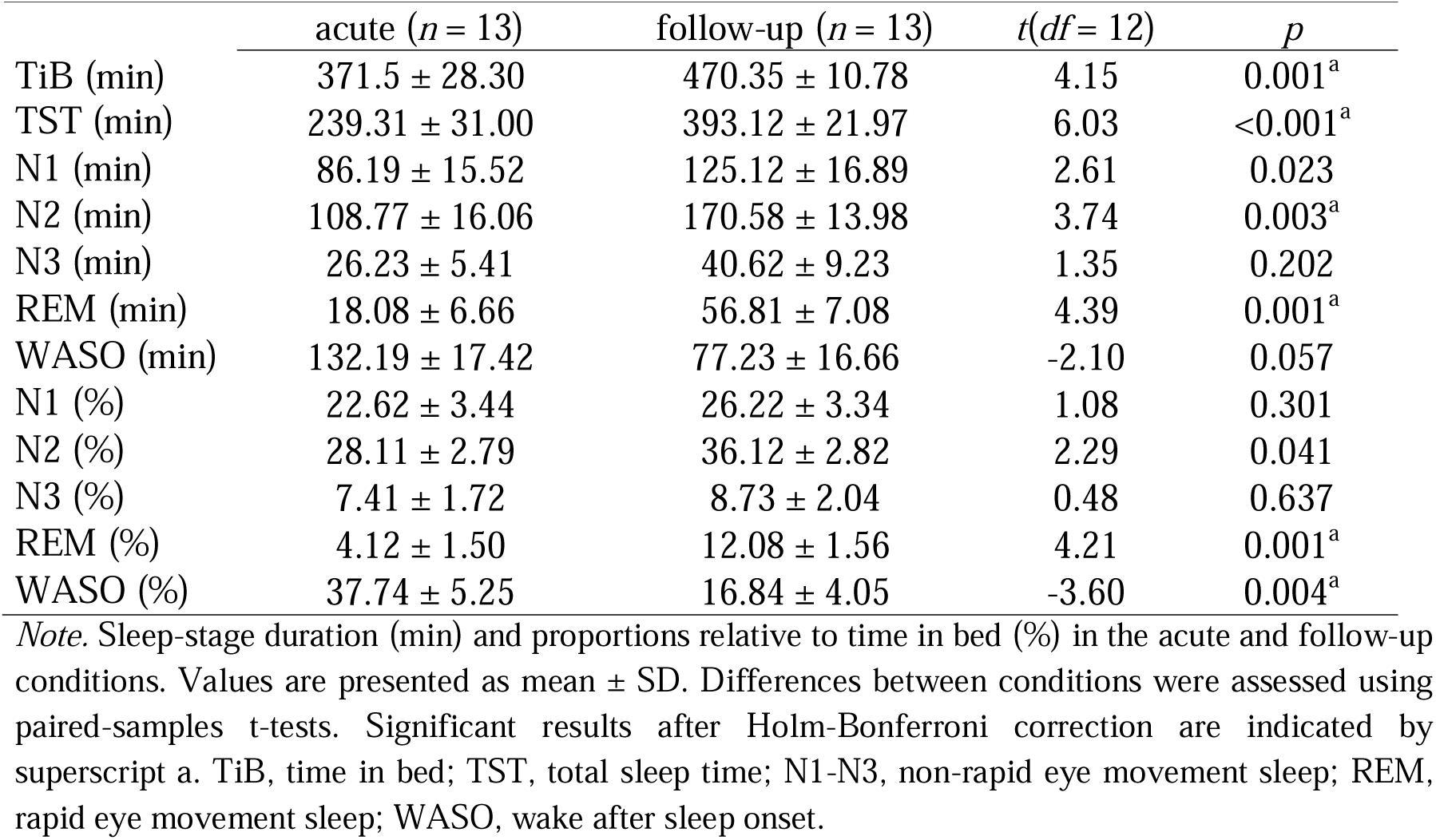
Sleep stage duration (minutes) and proportion of total sleep time (%) expressed as mean ± SEM during the acute and follow-up conditions.

Comparison of NREM power spectra showed significantly reduced theta activity during the acute phase compared to the follow-up recording (cluster-*p* = .014, Figure 3A).

**Figure 3.**
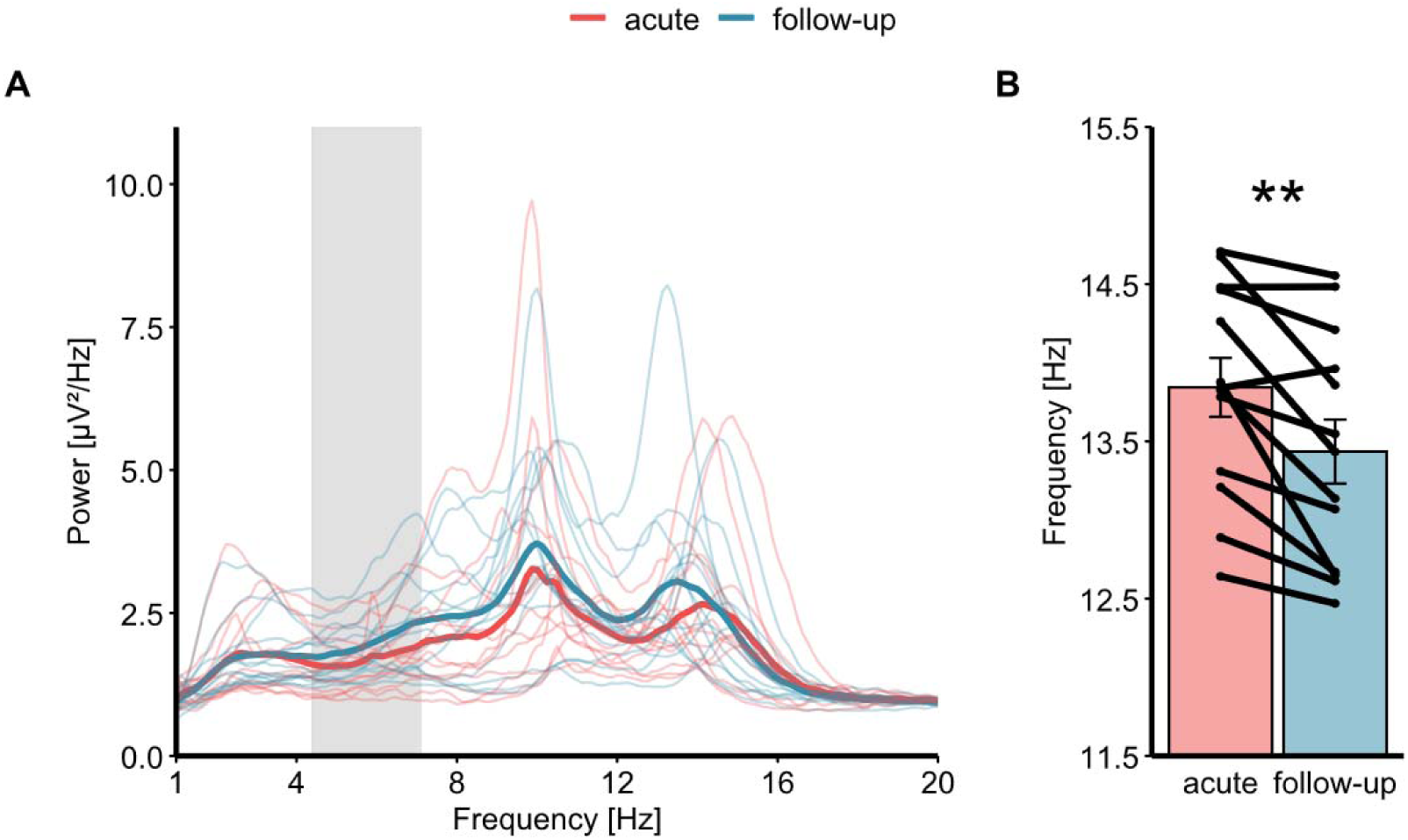
A). NREM power spectra (original minus aperiodic activity) during the acute and follow-up phase. The grey shaded area represents the significant cluster (cluster-*p* = .014) showing higher NREM theta power during follow-up than during the acute phase. B). Fast spindle frequencies during the acute and follow-up phase. \*\**p* < .01

NREM SO density and amplitude did not differ between the acute and follow-up sessions (all *p*’s > .63). Similarly, no significant differences were observed between conditions in density or amplitude of slow and fast spindles (all *p* > .70). Interestingly, spindle frequencies were higher during the acute phase compared to the follow-up phase (slow spindles: *t*(12) = -3,19, *p* = .008; fast spindles: *t*(12) = -3.82, *p* = .002; Figure 3B). Detailed spindle and SO characteristics are provided in Table S1.

To characterize SO-locked spectral dynamics, we first examined time-frequency representations (TFRs) time-locked to the SO trough in both phases. In both conditions, we observed increased activity in lower frequency ranges preceding the SO trough (all cluster *p*’s < .001) and increased fast spindle activity around the SO upstate (acute: cluster *p* < .001, follow-up: cluster *p* = .006; Figure 4). Participants did not differ significantly in their SO locked TFRs between conditions. Consistent with this finding, a comparable proportion of fast spindles co-occurred with an SO during the acute (7.90 ± 0.92%) and follow-up phase (8.50 ± 0.9 %; *t*(12) = -1.08, *p* = .300). Due to the overall reduced sleep duration during the acute state, the absolute number of spindle-SO co-occurring events was lower than during follow-up (*t*(12) = -3.71, *p* = .003). As a result, five participants had fewer than 20 co-occurring events during the acute phase and were thus excluded from analyses of preferred SO-spindle coupling phase. Among the remaining eight participants, spindle amplitude maxima were distributed non-uniformly across SO phases (Rayleigh test *p* < .01) in all but one participant during the follow-up session. In both the acute and follow-up conditions, the mean preferred coupling phase was oriented towards the SO upstate (0°, V-test *p* < .001). Moreover, the consistency of SO-spindle coupling did not differ between conditions (*t*(7) = - 0.84, *p* = .426).

**Figure 4.**
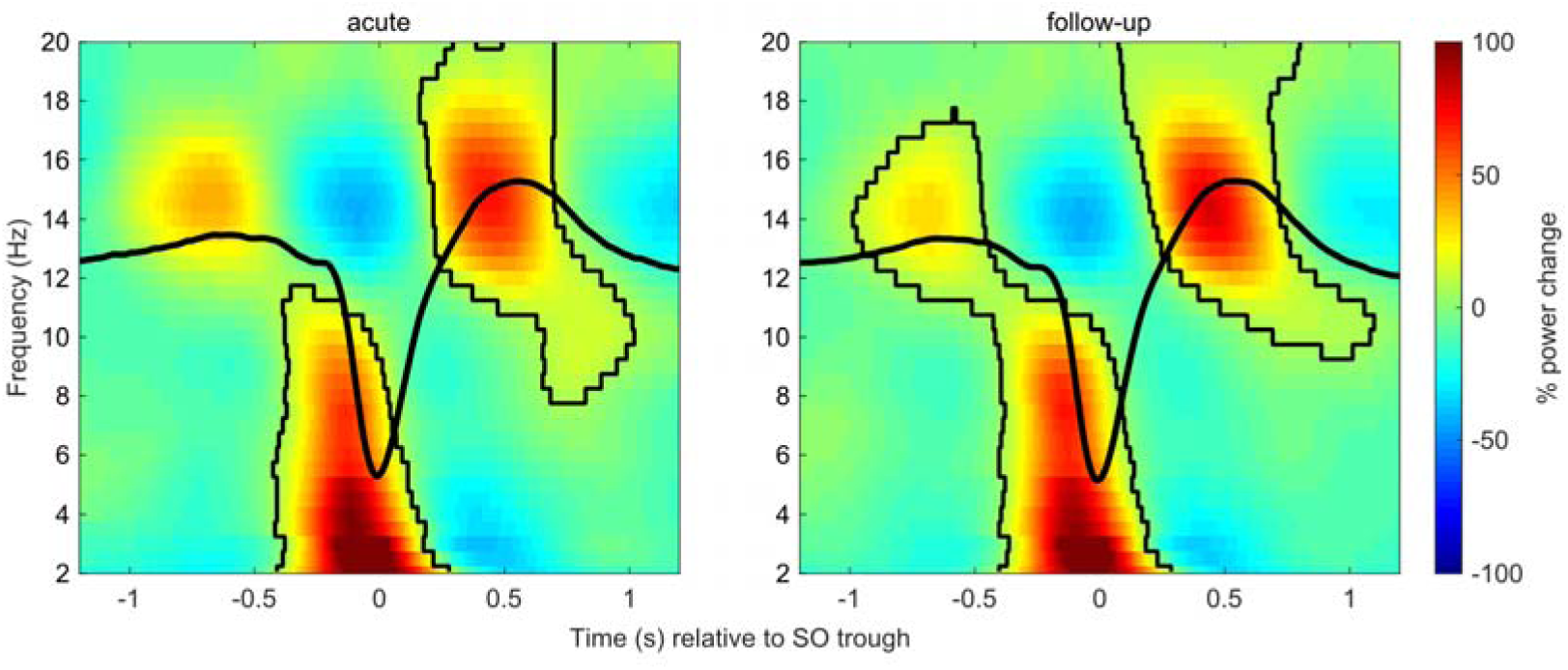
SO trough-locked time-frequency representations (TFRs) during the acute phase and the follow-up session, with the average SO waveform superimposed. Black contours indicate significant clusters of increased spectral power relative to baseline.

### Sleep oscillatory activity and memory consolidation

To examine brain-behavior associations, we fitted linear mixed-effects models with subject-specific random intercepts, including each sleep parameter, condition (acute vs follow-up), and their interaction as fixed effects. Given the observed difference in NREM theta power between the acute and follow-up recordings, and the proposed role of theta oscillations in hippocampal reactivation, we first tested whether aperiodic-corrected theta power predicted memory consolidation. The mixed-effects model revealed a significant theta power x condition interaction (β = 22.99, SE = 11.70, *z* = 1.97, *p* = .049, 95% CI [0.06, 45.92]; Figure 5A). In the acute condition, higher theta power tended to be associated with reduced overnight memory retention (β = -16.15, SE = 8.99, *z* = -1.80, *p* = .072, 95% CI [−33.75, 1.47]). The main effect of condition was not significant (β = -24.49, SE = 22.04, *z* = -1.11, *p* = .267, 95% CI [-67.69, 18.72]). To assess within-subject effects, we conducted a change-score analysis. Changes in theta power significantly predicted corresponding changes in memory consolidation (β = 53.00, SE = 23.38, *t*(11) = 2.27, *p* = .045, 95% CI [1.55, 104.46], *R*² = .318; Figure 5B).

**Figure 5.**
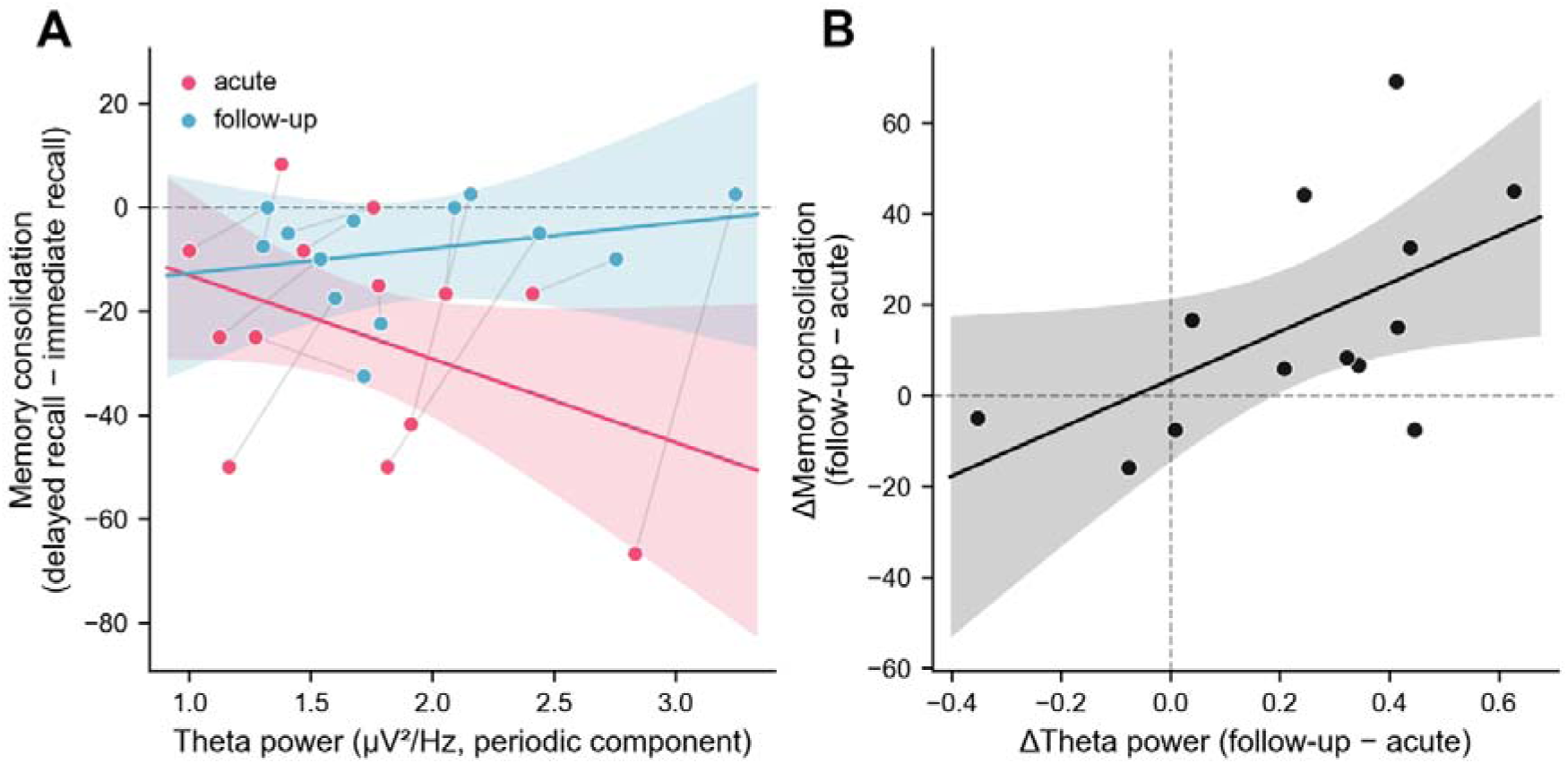
Theta power is differentially associated with memory consolidation across the acute and follow-up phases. A) Relationship between theta power and memory consolidation in the acute (red) and follow-up (blue) conditions. Solid lines represent linear regression fits, and shaded areas indicate the corresponding 95% confidence intervals. Gray lines connect individual participants across conditions. A significant theta power x condition interaction indicates that the association between theta power and memory consolidation differs between conditions. B) Within-subject association between changes in theta power and changes in memory consolidation from the acute to the follow-up condition. The solid line represents the linear regression fit, and the shaded area indicates the corresponding 95% confidence interval.

Fast and slow spindle density as well as SO density were examined in analogous exploratory models. For slow spindle density, the mixed-effects model showed neither a significant main effect (β = -7.20, SE = 16.14, *z* = -0.45, *p* = .655, 95% CI [-38.82, 24.42]) nor a significant interaction with condition (β = 11.23, SE = 23.82, *z* = 0.47, *p* = .637, 95% CI [-35.47, 57.92]). Likewise, the within-subject change analysis revealed no significant association with memory consolidation (β = -12.51, SE = 24.59, *t*(11) = -0.51, *p* = .621, *R*² = .023). For fast spindle density, the mixed-effects model did not yield a reliable interaction estimate due to convergence issues. Although the within-subject change analysis revealed a significant association between changes in fast spindle density and changes in memory consolidation (β = -54.98, SE = 23.99, *t*(11) = -2.29, *p* = .043, 95% CI [-107.79, -2.17], *R*² = .323), this finding should be interpreted cautiously because it was not supported by a corresponding interaction effect in the mixed-effects model. SO density did not significantly predict memory consolidation in the mixed-effects model (main effect: β = 0.042, SE = 0.029, *z* = 1.48, *p* = .140, 95% CI [-0.014, 0.098], interaction: β = -2.20, SE = 14.01, *z* = -0.16, *p* = .875, 95% CI [-29.66, 25.25]). Consistent with this result, the within-subject change analysis also showed no significant association (β = 0.036, SE = 0.042, *t*(11) = 0.86, *p* = .407, *R*² = .063).

To examine whether the association between theta activity and memory consolidation was independent of other sleep oscillations, we fitted a multivariable linear model including changes in theta power, SO density, and slow and fast spindle density. In this model, changes in theta power remained a significant predictor of memory consolidation (β = 67.59, SE = 28.58, *t*(8) = 2.36, *p* = .046). In contrast, changes in SO density (*p* = .084) and slow spindle density (p = .102) were not significantly associated with consolidation, whereas changes in fast spindle density showed a significant negative association (β = −59.92, SE = 24.64, *t*(8) = −2.43, *p* = 0.041).

Across oscillatory markers, theta power was the only measure that showed a consistent association with sleep-dependent memory consolidation across analytical approaches. Specifically, theta power was associated with consolidation both in the mixed-effects model and in the within-subject change analysis, and this relationship remained significant after controlling for other sleep oscillations. In contrast, classical NREM markers, including spindle and SO density, did not exhibit robust convergent associations. Although fast spindle density was significantly associated with memory consolidation in the multivariate model, this effect was not consistently supported across analyses and should therefore be interpreted cautiously.

## Discussion

Our findings show impaired overnight memory consolidation during the acute phase of TGA, a condition characterized by focal hippocampal CA1 lesions. Following clinical recovery, consolidation of verbal memory recovered, indicating restored sleep-associated memory formation. In parallel with these behavioral changes, sleep physiology differed between the acute TGA phase and follow-up. Specifically, NREM theta power was reduced during the acute phase of TGA and showed distinct associations with overnight memory consolidation across conditions. Importantly, increases in theta power from the acute to the follow-up phase predicted corresponding improvements in memory consolidation within individuals. In contrast, established NREM correlates of sleep-dependent memory consolidation, including sleep spindle density and SO density, did not differ between conditions and were not robustly associated with consolidation performance. Together, these findings suggest that transient CA1 dysfunction disrupts the sleep-dependent stabilization of newly encoded memories and is accompanied by a reduction in NREM theta activity, rather than by a global impairment of thalamocortical NREM sleep oscillations.

Consistent with our previous work, we observed pronounced deficits in episodic memory during the acute phase of TGA [35,33,36]. Across learning trials, patients showed significant improvements in performance in both phases, indicating that learning occurred during encoding despite overall reduced memory performance during the acute phase. This may reflect the fact that testing took place during later stages of the TGA episode, when patients had already begun to regain encoding capabilities [37]. Moreover, overnight retention was significantly lower during acute TGA compared with the follow-up session, indicating impaired sleep-dependent memory consolidation during transient hippocampal dysfunction. While previous studies in patients with TGA have demonstrated alterations in offline memory consolidation associated with transient CA1 dysfunction [33], the neurophysiological mechanisms underlying these changes have remained largely unexplored. The present study extends this work by linking sleep-dependent memory consolidation to specific sleep oscillations in patients with focal CA1 lesions. Because TGA is associated with transient focal dysfunction of the hippocampal CA1 region, these findings provide rare lesion-based evidence in humans that hippocampal CA1 integrity contributes to sleep-dependent memory consolidation.

Consolidation of episodic memories is thought to rely on a repeated reactivation of memory traces during sleep within hippocampal networks [3,38,1,39]. Thus, the deficits in overnight memory consolidation during the acute phase of transient CA1 dysfunction are consistent with a disruption of CA1-dependent processes involved in hippocampal replay and hippocampal-neocortical communication. Sharp-wave ripples and hippocampal replay cannot be measured directly using scalp EEG. Nevertheless, CA1 is a central component of hippocampal circuits involved in replay, synaptic plasticity, and output to downstream hippocampal and cortical targets [27,40,9,41]. The observed behavioral deficit therefore provides indirect evidence that transient CA1 dysfunction compromises the sleep-dependent stabilization of newly encoded memories.

Against this background, alterations in sleep-related oscillatory activity may provide indirect markers of disrupted hippocampal network dynamics during acute TGA. In support of this notion, another key finding of our study is the reduction of NREM theta power during the acute phase of TGA. Theta power increased following clinical recovery, suggesting that transient hippocampal dysfunction is associated with alterations in sleep-related theta activity. Importantly, the relationship between theta power and overnight memory consolidation differed between the acute and follow-up phase. Whereas higher theta power tended to be associated with greater overnight forgetting during the acute phase, higher theta power after recovery was linked to improved consolidation. Although the association observed during the acute phase did not reach conventional levels of statistical significance, the significant theta power x consolidation interaction indicates that the relationship between theta activity and memory consolidation depends on the state of the hippocampal network. Thus, theta activity may reflect distinct network states depending on the integrity of hippocampal CA1 circuits. Moreover, increases in theta power from the acute to the follow-up session predicted corresponding improvements in memory consolidation within individuals. Together, these findings indicate that changes in sleep-related theta activity are closely linked to behavioral recovery of memory consolidation following TGA. Importantly, theta power remained significantly associated with consolidation in a multivariable model controlling for SO density as well as slow and fast spindle density. Although fast spindle density showed a significant inverse association with consolidation in this exploratory analysis, this effect was not consistently supported across analytical approaches. By contrast, the association between theta power and memory consolidation was observed across multiple analyses, suggesting that NREM theta activity may represent a particular sensitive marker of hippocampal network integrity during sleep-dependent memory consolidation.

A central implication of these findings concerns current models of sleep-dependent memory consolidation. These models emphasize the hierarchical coordination of neocortical SOs, thalamocortical spindles, and hippocampal sharp-wave ripples as a mechanism for hippocampal-neocortical information transfer [12,5,13]. Our findings do not challenge this framework, but suggest that it may be incomplete without considering the contribution of theta activity. Converging evidence in humans indicates that theta activity may complement established sleep rhythms, including sleep spindles, in supporting hippocampal memory reactivation [16,23,15]. Theta activity has been shown to organize reactivation of hippocampus-dependent memory representations during both sleep and wakefulness, and these reactivation processes appear to share common neural signatures across physiological states [23]. Within this framework, reduced theta power during the acute phase of TGA may reflect impaired hippocampal network dynamics associated with transient CA1 dysfunction. Importantly, the altered relationship between theta activity and memory consolidation further suggests that theta power may reflect different functional network states depending on the integrity of hippocampal CA1 circuits. Thus, theta-related sleep activity may provide a complementary marker of hippocampal or hippocampal-neocortical network dynamics relevant for memory stabilization.

This interpretation is consistent with findings in Alzheimer’s disease and amnestic mild cognitive impairment, where alterations in sleep-related theta activity have been linked to disrupted memory-network function [42,43]. More broadly, Alzheimer’s disease is characterized by pronounced alterations in sleep physiology [44]. In our previous study of early Alzheimer’s disease, impaired sleep-dependent memory consolidation was related to changes in fast spindle amplitude, SO duration, and SO-spindle coupling [44]. Together with the present findings in TGA, these observations suggest that distinct forms of hippocampal network dysfunction may be reflected by different sleep-oscillatory profiles. Whereas transient CA1 dysfunction in TGA was primarily associated with reduced NREM theta activity, early Alzheimer’s disease was characterized by alterations in spindle- and SO-related dynamics, indicating that different pathophysiological mechanism may affect sleep-dependent memory processing through partially distinct oscillatory pathways.

Further support for this interpretation emerges from our recent work in patients with LGI1-associated limbic encephalitis, a condition affecting hippocampal subfields including DG-CA3 circuits [45]. In that study, impaired sleep-associated memory consolidation was accompanied by altered SO-spindle coupling despite largely preserved sleep macroarchitecture and comparable spindle and slow-oscillation densities. In contrast, the present findings in TGA show reduced NREM theta power during transient CA1 dysfunction whereas SO-spindle coupling remained largely preserved. Across these lesion models, the findings suggest that theta activity and SO-spindle coupling may reflect partially separable aspects of hippocampal-neocortical communication during sleep. More specifically, dysfunction of different hippocampal subfields may be associated with distinct alterations in sleep-oscillatory dynamics. However, this interpretation remains indirect and should be tested in future studies directly comparing hippocampal subfield dysfunction within a common experimental framework.

Thalamocortical spindles and SOs are widely considered as scalp EEG correlates of hippocampus-dependent memory consolidation [36,46,47]. In particular, sleep spindles have been proposed to facilitate the communication between hippocampal and neocortical memory systems by temporally coordinating hippocampal replay events with cortical plasticity processes [10,13,48]. Evidence from lesion studies further supports a link between hippocampal integrity, sleep oscillations, and memory consolidation. For example, patients with hippocampal sclerosis show alterations in spindle activity during slow-wave sleep, and spindle density predicts the degree of sleep-related memory strengthening [49]. Similarly, altered SO-spindle coordination has been reported in patients with bilateral hippocampal damage [50]. In the present study, however, spindle density and amplitude did not differ between the acute and follow-up phase, and spindle measures were not robustly associated with memory consolidation. These findings suggest that the basic generation of thalamocortical spindle activity remains largely preserved during transient hippocampal CA1 dysfunction. Likewise, the temporal coupling of spindles to the SO upstate did not differ between conditions, indicating that large-scale coordination of thalamocortical sleep oscillations remains intact despite focal hippocampal lesions. Interestingly, spindle frequencies were slightly higher during the acute phase than during the follow-up recording. Changes in spindle frequency have previously been linked to alterations in thalamocortical network dynamics and neuromodulatory states during sleep [51,52]. Together with the observed alterations in theta activity, these findings suggest a dissociation between theta-related and thalamocortical sleep dynamics. Transient CA1 dysfunction was associated with reduced NREM theta power, whereas the generation and temporal coordination of thalamocortical SO and spindles were largely unaffected. This pattern argues against a non-specific disturbance of NREM sleep physiology and instead points to theta activity as the NREM oscillatory measure most closely tracking memory recovery in the present sample.

Several limitations should be considered when interpreting the present findings. First, the sample size was relatively small, reflecting both the rarity of TGA and the practical challenges of obtaining polysomnographic recordings during the acute state of the syndrome. Replication in larger samples will therefore be important to confirm the observed associations between sleep oscillations and memory consolidation. Second, the present study relied on scalp EEG recordings, which do not allow direct assessment of hippocampal activity such as SWRs. Consequently, the interpretation of sleep-related theta oscillations as an indirect marker of hippocampal network dynamics remains inferential. Third, sleep architecture differed between the acute and follow-up recordings, including reduced total sleep time and REM sleep during the acute phase. Although slow-wave sleep did not differ significantly between conditions, we cannot fully exclude the possibility that changes in sleep macroarchitecture contributed to the observed alterations in memory consolidation. Finally, the associations between sleep oscillations and behavioral consolidation were correlational and therefore do not allow causal inferences regarding the underlying mechanisms.

Future research should extend these findings by directly testing whether theta activity plays a causal role in sleep-dependent memory reactivation or instead reflects the functional state of hippocampal-neocortical networks. One promising approach would be to combine targeted memory reactivation with high-density EEG, MEG, or intracranial recordings in patients with focal hippocampal dysfunction, to determine whether experimentally cued reactivation elicits theta responses that predict subsequent memory retention [18,23,53,54]. Another important step will be to test whether non-invasive stimulation protocols designed to modulate theta activity during NREM sleep can influence memory consolidation, building on previous stimulation approaches that have successfully altered sleep oscillations and memory processing [54]. Such studies would help determine whether theta activity is merely a marker of hippocampal network integrity or contributes mechanistically to the stabilization of newly encoded memories during sleep.

Overall, our findings highlight the importance of hippocampal CA1 integrity for sleep-dependent memory consolidation in humans. While classical thalamocortical sleep oscillations such as SOs and sleep spindles remained largely preserved during the acute phase of TGA, NREM theta activity was reduced and closely linked to changes in memory consolidation. These findings suggest that sleep-related theta oscillations may provide an indirect marker of hippocampal-neocortical network function during memory consolidation, including compensatory reorganization during recovery. By showing that memory impairment and theta alterations recover in parallel following TGA, the present study extends current models of sleep-dependent consolidation and highlights theta activity as a promising marker of hippocampal network integrity and a potential target for future mechanistic investigations.

## Methods

### Patient cohort

Fourteen patients (mean age 65.79 ± 5.81 years; range 55 – 75 years; 9 female) who presented to our neurological emergency unit during the acute phase of a TGA participated in the study. The amnesic syndrome was diagnosed according to established diagnostic criteria for TGA [35,55], including: i) an attack witnessed by an observer present for most of the attack, ii) a clear anterograde amnesia during the attack, iii) no clouding of consciousness or loss of personal identity, vi) cognitive impairment limited to amnesia, v) absence of focal neurological symptoms or epileptic signs, vi) no recent history of head injuries or seizures, and vii) resolution of symptoms within 24 hours. The characteristic clinical course of TGA includes an abrupt onset of severe hippocampus-dependent memory impairment followed by gradual recovery of memory function during the last third of the attack. To facilitate rapid patient recruitment, a neurologist was available on-call throughout the study period for patient evaluation. A follow-up measurement took place 8.5 ± 4.2 months (range:1.73 – 32.73 months) after the acute phase of the TGA. Both the acute and follow-up assessments included a standard neurological examination. Patients also underwent a structured interview to assess the temporal course of the TGA episode, relevant clinical factors, and a history of cardiovascular and neurological disease. All patients provided written informed consent prior to participation. The study was approved by the Ethical Committee of the University of Kiel and was conducted in accordance with the Declaration of Helsinki.

### Materials and procedure

Declarative memory was assessed using a paired-associate learning task administered before and after a night of sleep in both the acute and follow-up conditions. Testing during the acute phase was conducted during inpatient treatment at the Department of Neurology at the University Hospital Kiel. Follow-up testing was conducted on an outpatient basis in the sleep laboratory. The task consisted of 12 German word pairs in the acute condition and 40 pairs in the follow-up condition. During acquisition, patients were instructed to generate a sentence containing both words of each semantically associated word pair (e.g., furniture - chair). This was followed by a period of quite reading of the complete list lasting 6 minutes in the acute phase and 15 minutes in the follow-up condition. Immediate recall was assessed 30 minutes after encoding. Patients were presented with the first word of each pair and were asked to recall the corresponding target [56–58]. Responses were recorded by the experimenter, and the correct target word was provided irrespective of response accuracy. The study-test cycle was repeated five learning trials. Participants were allowed to sleep from 11:00 p.m. onward. Delayed recall was assessed after an approximately 10-hour retention interval that included a full night of sleep and was administered between 9:00 and 10:00 a.m the following morning.

### Polysomnographic recording and EEG analysis

Polysomnographic recordings included electroencephalography (EEG) from electrodes F3, F4, C3, C4, O1, and O2 according to the international 10-20 system, referenced online to Cz. Bilateral mastoid electrodes were recorded to permit offline re-referencing to the average mastoid signal. In addition, electrooculography (EOG), electromyography (EMG), and electrocardiography (ECG) were recorded. EEG and EOG signals were band-pass filtered between 0.2 and 35.0 Hz. EMG signals were filtered between 2.0 and 128.0 Hz, and ECG signals between 0.5 and 1000 Hz. Data were acquired using a SOMNOscreen^TM^ plus system (Somnomedics, Randersacker, Germany) at sampling rates of 128 or 256 Hz. To ensure consistency across recordings, all data were down-sampled to 128 Hz prior to analysis. Sleep stages were scored according to the criteria of the American Academy of Sleep Medicine (AASM) [59] by a trained rater. Total sleep time (TST), wake after sleep onset (WASO), time spent in stage N1, stage N2, slow wave sleep (SWS/N3), and REM sleep were quantified in 30–s epochs. Artifacts identified during sleep scoring were excluded from all further analyses. All EEG analyses were restricted to central channels because frontal derivations were unavailable in some recordings and occipital areas generally exhibited poor signal quality. In all but six recordings, for which only either C3 or C4 met quality criteria, EEG measures were calculated from the average of C3 and C4. One participant was excluded due to poor EEG signal quality during recording in the acute phase, resulting in a final sample of 13 participants with both acute and follow-up polysomnographic recordings available for analyses.

NREM power spectral density was calculated in MATLAB R2023a from artefact-free 5s segments using Welch’s method (MATLAB function *pwelch*; 50% overlap, Hamming window). Power spectra were subsequently analyzed using the Python-based toolbox “fitting oscillations and one-over f” (fooof) to separate periodic and aperiodic spectral components [60]. Spectral parametrization was performed in a frequency range from 0.5 to 25 Hz using the fixed mode. After removal of the aperiodic component (1/*f* like activity), fitted peaks within the slow-spindle (acute: 10.46 ± 0.16 Hz; follow-up: 10.21 ± 0.10 Hz) and fast-spindle range (acute: 14.29 ± 0.19 Hz, follow-up: 13.82 ± 0.19 Hz) were visually inspected and manually adjusted when necessary. These individual peak frequencies served as center frequencies for the detection of spindles. Two participants did not exhibit a detectable slow-spindle frequency peak; in these cases the mean slow-spindle peak frequency of the respective condition was used for spindle detection. Sleep spindle and SO-oscillation detection was performed on artefact-free NREM epochs using the SleepTrip toolbox (RRID: SCR_017318) with parameters described previously [61]. For both spindles and SOs, event density (events/minute) and amplitude (µV; trough to peak potential) were quantified. For spindles, mean oscillatory frequency (Hz) was additionally determined.

Time-frequency representations (TFRs) of SOs were computed for epochs extending ± 3s around each SO trough using the FieldTrip toolbox in MATLAB [62]. Spectral decomposition was performed using Morlet wavelets with a linearly increasing number of cycles (4 to 12 cycles) across frequencies from 2 to 20 Hz in steps of 0.5 Hz. For each participant and channel, power was normalized as the percentage change relative to a baseline period defined as the mean power within ± 1.5s around the SO trough. Spindle-SO co-occurrence was evaluated by determining, for each detected spindle, whether it occurred between the two positive-to-negative zero crossings of an SO. The proportion of spindles co-occurring with a SO was then calculated. As a last step, we calculated the phase-amplitude coupling of SOs and fast spindles. Therefore, the EEG signal was filtered in the SO (0.3 Hz 4^th^ order twopass high-pass and 4 Hz, 6^th^ order, twopass low-pass Butterworth filter) and the spindle frequency band (± 1.5 Hz around the individually determined fast spindle frequency peak in the power spectrum; 4^th^ order, twopass Butterworth filter). For each spindle occurring within the two positive-to-negative zero crossings of a SO, we extracted the instantaneous phase of the SO–filtered signal at the time point corresponding to the maximum amplitude of the spindle filtered signal. Circular means and vector lengths were calculated using the CircStat toolbox in MATLAB [63].

### Magnetic resonance imaging

Clinical whole-brain MRIs were acquired 24-72 h after onset of TGA symptoms when the detectability of hippocampal lesions is highest [29]. High-resolution MRI was acquired on a 3 Tesla scanner (Philips Achieva) using diffusion-weighted Echo Planar Imaging (DW-EPI; TR/TE/FA 319/2.4/80, slice thickness 2mm, voxel size 1.67 x 2.12 x 3 mm) with subsequent maps of the apparent diffusion coefficient (ADC), as well as additional T2-weighted turbo spin echo sequences (TR/TE/FA = 4025/100/90, slice thickness 2mm, voxel size 0.51 x 0.65 x 2 mm) transverse oblique plane parallel to the hippocampus and coronal perpendicular to the hippocampus. The whole brain including temporal and frontal lobe structures was inspected with respect to structural abnormalities.

Lesions were classified as hippocampal CA1 lesions only when detectable in both DWI and T2-weighted images with hyperintense DWI and T2 lesions corresponding to identical locations within the different sectors of the cornu ammonis in the coronal plane and the rostral-occipital position within the hippocampus. MR images were visually inspected by two neuroradiologists and one neurologist with extensive experience in the detection of deviations in hippocampal signals in diagnostic investigation of TGA. Lesions were mapped within the different sectors of the cornu ammonis after Lorente de Nó according to the anatomical reference atlas of Duvernoy [64].

### Neuropsychological assessment

At follow-up participants completed a neuropsychological test battery after delayed recall of the paired-associate learning task on the morning following the sleep recording. The battery comprised the Rey-Osterrieth Complex Figure Test, Trail Making Test parts A and B, the Regensburg Word Fluency Test, and a German version of the National Adult Reading Test (Mehrfachwahl-Wortschatz-Intelligenztest-B). The assessment was used to characterize cognitive functions in domains including visuospatial memory, attention, processing speed, executive function, verbal fluency, and premorbid general intelligence. Patients performed within a normal range when recovered from the amnesic phase (Table 2).

**Table 2.** Neuropsychological performance at follow-up (n = 14). Values are presented as mean ± SEM.

| | Mean ± SEM |
| --- | --- |
| ROCF copy | 32.69 ± 0.47 |
| ROCF recall | 17.69 ± 1.12 |
| RWT (letter S) | 17.43 ± 1.18 |
| RWT (letter P) | 13.79 ± 1.29 |
| TMT-A | 45.92 ± 4.50 |
| TMT-B | 100.43 ± 7.50 |
| MWT-B | 32.36 ± 0.94 |

Note. ROCF = Rey-Osterrieth complex figure task, RWT = Regensburg word fluency task, TMT = trail making test, MWT-B = Mehrfachwahl-Wortschatz-Intelligenztest-B (German version of the National Adult Reading test)

### Analysis of cortisol

As stress and elevated cortisol levels have been found to modulate sleep-dependent memory consolidation [65,66], salivary cortisol levels were assessed in all participants. Saliva samples were collected in late evening prior to memory testing and immediately stored at -25°C until analysis via immunoassay (Immulite, Siemens Healtcare, Erlangen, Germany, intra-assay coefficient of variation 6.8%, inter-assay coefficient of variation 9.9%). Mean cortisol levels during the acute phase of TGA were 1.1 ± 0.41 ng/ml (range 0.1 – 3.3 ng/ml, normal circadian matched range 0.04 – 4.67 ng/ml) and consistent with normal nocturnal suppression of cortisol secretion. One patient showing significantly elevated cortisol levels was excluded from further analyses.

### Statistical analyses

Statistical analyses were conducted using IBM SPSS Statistics (Version 29), R Studio (version 4.4.2), and Python (Version 3.13). Behavioral performance across learning trials and retention intervals was analyzed using repeated-measures ANOVAs. Memory consolidation was quantified as the difference between delayed recall after sleep and immediate recall before sleep (consolidation index = delayed recall - immediate recall). Immediate recall was defined as the highest level of performance achieved across the five learning trials prior to sleep, reflecting the maximal degree of encoding attained before the retention interval. Significant interaction effects in ANOVA models were followed up by paired-sample t-tests. The significance level was set at p < .05 (two-tailed). Where appropriate, Holm-Bonferroni correction was applied to control for multiple comparisons. Data are reported as mean ± SEM.

Power spectra (original spectra minus aperiodic activity) during the acute and follow-up phase were compared using paired-samples t-tests, correcting for multiple comparisons using cluster-based permutation testing [67] with 5000 permutations and a cluster-forming threshold of p < .05 (two-tailed). The baseline-normalized TFRs were tested against zero for each condition, similarly, using cluster-based permutation tests to correct for multiple comparisons. Using paired-samples t-tests the baseline-normalized TFRs at both conditions were then compared and corrected for multiple comparisons using cluster-based permutation tests. Distribution of preferred phases of SO-spindle coupling were tested using the Rayleigh and V-test in the CircStat toolbox [63].

Associations between sleep oscillatory activity and memory consolidation were examined using linear mixed-effects models estimated with restricted maximum likelihood and subject-specific random intercepts. Based on our a priori hypothesis regarding the role of theta oscillations in memory reactivation during sleep, theta power was specified as the primary predictor. In addition, established NREM sleep markers implicated in memory consolidation, including fast spindle density, slow spindle density, and SO density, were examined in secondary analyses. Fixed effects included condition (acute vs. follow-up), the respective sleep parameter, and their interaction. If the variance component of the random intercept was estimated to be negligible, indicating minimal between-subject variability, complementary within-subject change-score analyses were conducted as a sensitivity analysis to assess the robustness of state-dependent associations between sleep parameters and memory consolidation. To further examine whether the association between theta activity and memory consolidation was independent of other sleep oscillations, a multivariable linear regression model was fitted including changes in theta power, SO density, and slow and fast spindle density as simultaneous predictors. Model assumptions were evaluated by visual inspection of residual distributions and residual-versus-fitted plots. Potential influential observations were examined using Cook’s distance.

## Acknowledgements

Funding: This study has been supported by the German Research Foundation (DFG) FOR 5434, by the Else Kröner-Fresenius-Stiftung and by the Faculty of Medicine, University of Kiel, Germany.

## Author contributions

T.B., A.H., and J.B. designed the study. A.H. and E.M.K. performed the analyses, interpreted the data, and drafted the manuscript. F.D.W. contributed to data analysis and interpretation. J.D. collected the data. A.B. and A.P. provided supervision and critically revised the manuscript. All authors contributed to the final version of the manuscript and approved its submission.

## Competing interests

The authors declare no competing interests.

## Data Availability Statement

The datasets generated and analyzed during the current study are available from the corresponding author on reasonable request.

## Notes

### Competing Interest Statement

The authors have declared no competing interest.

