## Supplemental Table 1 for "Hippocampal CA1 neurons are crucial for sleep-associated memory formation in humans: The role of theta power during NREM sleep"

**Table S1. Slow Oscillation and spindle characteristics (M ± SEM)**

|  | acute (*n* = 13) | follow-up (*n* = 13) | *t*(*df* = 12) | *p* |
| --- | --- | --- | --- | --- |
| Slow Oscillations |  |  |  |  |
| density (#/min) | 2.834 ± 0.293 | 2.857 ± 0.23 | 0.14 | 0.893 |
| amplitude (µV) | 115.814 ± 7.413 | 117.944 ± 7.585 | 0.49 | 0.630 |
| Slow Spindles |  |  |  |  |
| density (#/min) | 3.392 ± 0.179 | 3.369 ± 0.163 | -0.14 | 0.894 |
| amplitude (µV) | 26.957 ± 2.253 | 26.63 ± 2.042 | -0.25 | 0.806 |
| frequency (Hz) | 10.165 ± 0.15 | 9.881 ± 0.11 | -3.19 | 0.008 |
| Fast Spindles |  |  |  |  |
| density (#/min) | 4.109 ± 0.199 | 4.164 ± 0.161 | 0.39 | 0.707 |
| amplitude (µV) | 21.705 ± 1.537 | 21.766 ± 1.431 | 0.07 | 0.943 |
| frequency (Hz) | 13.844 ± 0.187 | 13.435 ± 0.204 | -3.82 | 0.002^a^ |

The significance of differences between acute and follow-up are tested using paired samples t-test.
^a^ Significant after Holm-Bonferroni correction.
